# The cap size and shape of *Arabidopsis thaliana* primary roots impact the root responses to an increase in medium strength

**DOI:** 10.1101/378828

**Authors:** J. Roué, H. Chauvet, N. Brunel-Michac, F. Bizet, B. Moulia, E. Badel, V. Legué

## Abstract

During root progression in soil, root cap cells are the first to encounter obstacles. The root cap is known to sense environmental cues, making it a relevant candidate for a mechanosensing site. An original two-layer medium was developed in order to study root responses to growth medium strength and the importance of the root cap in the establishment of these responses. Root growth and trajectory of primary roots of *Arabidopsis thaliana* seedlings were investigated using *in vivo* image analysis. After contact with the harder layer, the root either penetrated it or underwent rapid curvature, enabling reorientation of the root primary growth. The role of the root cap in tip reorientation was investigated by analyzing the responses of *Arabidopsis* mutant roots with altered caps. The primary root of *fez-2* mutant lines, which has fewer root cap cell layers than wild-type roots, showed impaired penetration ability. Conversely, *smb-3* roots of mutant lines, which display a higher number of root cap cells, showed enhanced penetration abilities. This work highlights that alterations in root cap shape and size affect the root responses to medium strength.

**Highlight:** The analysis of the growth and orientation of *Arabidopsis thaliana* mutant roots affected in root cap size and shape showed that properly formed root cap is required to trigger the root responses to medium strength.

**Abbreviations:**

COL: columella;
LRC: Lateral Root Cap;
SI: Sharpness Index;
SMB: SOMBRERO.

## Introduction

Roots grow in a complex soil environment, which exhibits spatial and temporal heterogeneities, arising from sand particles, stones or hardpans of soil that constitute physical obstacles. The ability of roots to penetrate the soil and to overcome physical obstacles depends either on their ability to find a path of least resistance, or on their ability to generate sufficient force to go through the obstacle (Bengough *et al*., 2009; Vollsnes *et al*., 2010; Colombi *et al*., 2017*a*). The force needed to penetrate soil, provided by root growth, is determined by soil strength. In natural conditions, soil strength – commonly estimated by penetrometer measurements – is in the range of 1MPa (Jin *et al*., 2013; Kolb *et al*., 2017). Soil heterogeneities may be overcome with soil compaction or soil drying, which results in an increase in soil strength.

In a few studies carried out in laboratories, changes in soil strength have been mimicked with simplified models adapted to the study of root responses. The elaboration of a two-layer Phytagel-based growth medium, containing two layers of distinct Phytagel concentrations, enables the roots to experience variations in medium strength when penetrating from one medium layer to another (Yamamoto *et al*., 2008; Yan *et al*., 2017). It has been shown that adjusting the Phytagel concentration modulated growth medium strength (Schiavi *et al*., 2016). In two-layer media experiments, where roots faced a compliant layer and a stiff layer consecutively, root tips were submitted to increasing axial stress as they encountered the interface between the two layers. Roots which did not manage to generate sufficient force to penetrate the lower layer underwent buckling, dramatically changing the root growth trajectory and the root straightness (Massa and Gilroy, 2003; Silverberg *et al*., 2012; Bizet *et al*., 2016). The maximal axial force a root is able to exert before buckling, given by Euler’s law, depends on the rigidity, the length of the growth zone, and the diameter of the root to the power of four (Timoshenko and Goodier, 1970). In accordance with Euler’s law, species with a higher increase in root diameter in response to medium strength are more efficient in penetrating strong layers (Materechera *et al*., 1992). Besides root diameter and growth zone length, it has been shown that root tip shape was also an important feature modulating root responses to medium strength (Ruiz *et al*., 2016; Colombi *et al*., 2017*b*). In compacted soils, root tips with an acute opening angle grow faster than root tips with a blunted geometry (Iijima *et al*., 2003*a*,*b*; Colombi *et al*., 2017*b*). Colombi et *al*. (2017) showed that root tip shape influenced the penetration stress experienced by roots, by modulating distribution of the local compaction concentrated around the tip.

The shape of root tips arises from the shape of the root cap that covers and protects the root apical meristem. The root cap is composed of the Columella (COL) located at the centre of the root cap and the Lateral Root Cap (LRC) (Arnaud *et al*., 2010; Kumpf and Nowack, 2015). In *Arabidopsis thaliana*, the mature COL and LRC are organized in 3 to 5 cell layers of increasing age. Every formative division of root cap stem cells generates a new differentiating cell layer which pushes the older root cap layers towards the root periphery. COL mature cells are known to sense gravity, thanks to the amyloplasts they contain, conferring to the root cap the ability to trigger tropisms in the root (Sato *et al*., 2015). LRC mature cells principally act as secretory cells which synthesize and export high molecular weight polysaccharide mucilage (Kumpf and Nowack, 2015). Once mature, COL and LRC cell layers reach the periphery of the root cap, where they end up being sloughed off, allowing the root cap to keep a constant size (Barlow, 2003). The balance between cell production and cell sloughing controls the changes in root tip shape. The study of root cap formation provided for the identification of the molecular factors involved in cell production, differentiation and sloughing. Among these factors, FEZ and SOMBRERO (SMB) transcription factors from the NAC family [NAM (NO APICAL MERISTEM), ATAF (*ARABIDOPSIS* TRANSCRIPTION ACTIVATION FACTOR) and CUC (CUP-SHAPED COTYLEDON)] were identified. FEZ specifically promotes periclinal divisions of COL and LRC stem cells (Willemsen *et al*., 2008). The *fez-2* mutant, resulting from the loss of function of *FEZ*, exhibits a lower rate of periclinal division of the root cap stem cells which leads to the reduction of COL and LRC cell layers compared to wild-type roots (Willemsen *et al*., 2008; Bennett *et al*., 2014). SMB prevents the division of the differentiating root cap daughter cells and promotes their differentiation. Additional COL and LRC cell layers are observed in the *smb-3* loss-of-function mutant and the outer LRC cell layer reaches the end of the elongation zone (Willemsen *et al*., 2008; Bennett *et al*., 2010).

The objective of our study is to investigate the role of the structure of the root cap in the establishment of root responses to medium strength. We hypothesized that pointed caps facilitated root penetration and blunted or domed caps complicated root penetration. We studied the root response to medium strength through the *fez-2* and *smb-3 Arabidopsis thaliana* mutant lines because of their contrasted cap structure. We characterized the root tip morphology of *fez-2* and *smb-3* mutants to understand whether the loss or the gain of root cap cell layers impacted root tip size and shape. Then, Col-0, *fez-2* and *smb-3* roots were grown in two-layer Phytagel media of increasing mechanical resistance. The mutants were studied using a novel experimental set-up that enabled quantitative imaging of the growth and orientation of the root tip. Using a spatiotemporal analysis, we characterized the ability of the root to penetrate the harder medium, the root growth rate, and the root curvature of wild-type and mutant roots. We discovered that penetration ability and the reorientation were affected in *fez-2* and *smb-3* primary roots in exactly the opposite way to what was hypothesized.

## Material and Methods

### Plant material and growth conditions

The wild-type strain of *Arabidopsis thaliana* (ecotype Columbia, Col-0) and *fez-2, smb-3 Arabidopsis thaliana* null-mutants used for the experiments were provided by B. Scheres’s laboratory (Willemsen *et al*., 2008; Bennett *et al*., 2010, 2014). For all the experiments, Phytagel-based growth media were elaborated. We used Phytagel as a substitute for Agar for its higher temperature stability, higher mechanical strength and better optical clarity (Schiavi *et al*., 2016). The growth medium consisted of half-strength Murashige and Skoog (MS/2) medium adjusted to pH 5.7 and containing Phytagel at different concentrations. Two-layer media were developed, based on the superposition of an upper layer of growth medium containing 0.2% (w/v) of Phytagel and a lower layer containing 0.2 % (w/v), 0.3% (w/v) or 0.5 % (w/v) of Phytagel (Fig.S1). According to these Phytagel compositions, the two layer-media were called 0.2/0.2, 0.2/0.3 and 0.2/0.5. The two-layer media were poured into specific culture chambers which consisted of polycarbonate boxes measuring 100 X 20 mm, and which were 60 mm deep, allowing roots to grow vertically inside the growth medium. The two-layer media were prepared as follows: 60 mL of sterilized MS/2 medium containing 0.2% (w/v), 0.3% (w/v) or 0.5% (w/v) of Phytagel were poured into the culture chambers under aseptic conditions. These 60 ml formed the 30 mm deep lower medium layer. After 10 min, 20 mL of a sterilized MS/2 medium containing 0.2% (w/v) of Phytagel were poured over the lower medium layer and formed the 10 mm deep upper layer.

After the two-layer media preparation, the Arabidopsis seeds were surface sterilized with 30% (v/v) sodium hypochlorite solution, added to 2% (v/v) Triton X-100 for 10 min. After 5 washes in sterile distilled water, 10 seeds were gently placed on the surface of the growth medium. After sowing, the culture chambers were sealed and kept at 4 °C in the dark for 2 days. They were then placed at 23 °C under a 16 h-photoperiod.

### Mechanical properties of Phytagel growth medium

We performed compressive and penetration tests on each Phytagel medium, in order to determine their elastic modulus and mechanical penetration resistance.

#### Elastic modulus measurement

Growth media containing from 0.2% to 1.2% (w/v) of Phytagel were poured into 10 mm thick Petri dishes. Then, 20×20×10 mm^3^ Phytagel samples were cut and placed in a universal testing device (Instron 5565). A uniaxial compressive load was applied on the surface of the gels by displacement of the upper plate with a constant loading rate of 0.5 mm.min^-1^ (Fig. 1A). A 100 N load cell recorded the force needed to deform the samples. After a typical contact step, the loading force increased with compression of the sample, before decreasing notably with the slipping or the rupture of the sample. Real-time load and displacement recordings provided the characteristic mechanical stress-strain curves (Fig. 1B). In the linear portion of the curve, *i.e*. when stress is proportional to strain, deformation is considered to be elastic (Schiavi *et al*., 2016). In this range, the elastic modulus (E) value of the Phytagel samples was determined from the ratio between the incremental stress Δσ and the incremental strain rate Δε according to Hooke’s law:

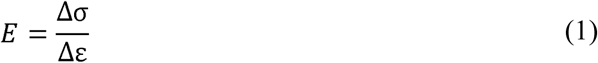

**Fig. 1.**
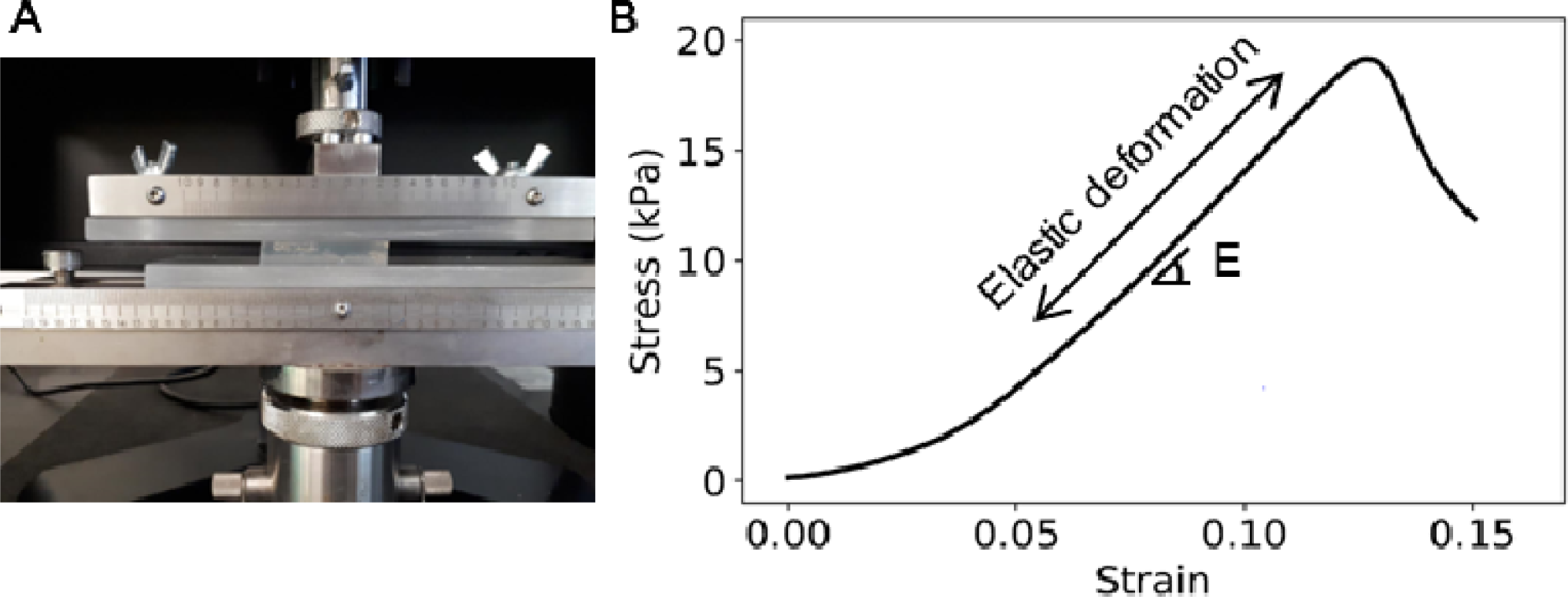
Characteristic stress-strain curve obtained *via* compressive tests. (A) Representation of 1.2% Phytagel cube placed on Instron device and compressed by an upper plate at a loading rate of 0.5mm/min. (B) Stress-strain curve resulting from a compressive test performed on a 1.2% Phytagel medium sample. The Elastic modulus E corresponds to the slope of the linear portion of the curve, which is the zone of elastic deformation.

Where

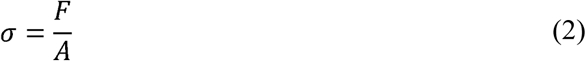

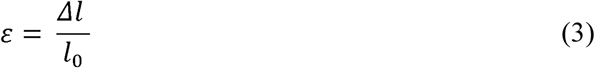

and *F* is the compression force (N), *A* is the surface area of the cross-section of the sample (m^2^) on which the force is applied, *l*_*0*_ is the initial thickness of the sample (m), Δ*l* is the variation of this thickness, *i.e*. displacement of the plate (m). Compressive tests were repeated for 3 samples of each Phytagel concentration.

#### Penetration resistance measurement

Penetration tests were conducted in 0.2/0.2, 0.2/0.3 and 0.2/0.5 two-layer media. The Phytagel medium samples were prepared in the same culture chambers used for root growth. A 2.5 mm diameter needle was attached to a load cell (Instron, 100 N max) and moved downward into the gel at a rate of 1 mm.min^-1^. Real-time recording of the force-displacement curve made it possible to determine the maximal force required for the rupture of the interface between the upper and the lower layer.

### Root tip shape and determination of Sharpness Index

5-day-old primary roots of the *Arabidopsis thaliana* Col-0, *fez-2* and *smb-3* lines, growing in a 0.2% one-layer medium were extracted from the medium and placed on a microscope slide. The root tips were stained with 10 µg.ml^-1^ propidium iodide (PI, Sigma Aldrich, St-Louis, MO, USA) and observed with an Apochromat X20 lens coupled to a Zeiss LSM 800 laser scanning microscope (Zeiss, Jena, Germany).

To characterize the root tip size and shape of each genotype, the longitudinal median section from the quiescent centre to the root tip were extracted from longitudinal root observations (Fig. 2). The length of the vertical section *L*_*R*_, the diameter of the root at the quiescent centre *D*_*R*_ and the area of the radial section of the root tip *A*_*R*_ were measured using ImageJ software (v1.52d, http://imagej.nih.gov/ij/) (Fig. 2). An *inverse* Sharpness Index (*i*SI) of the root tip was calculated according to the following equation:

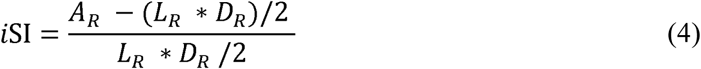

**Fig. 2.**
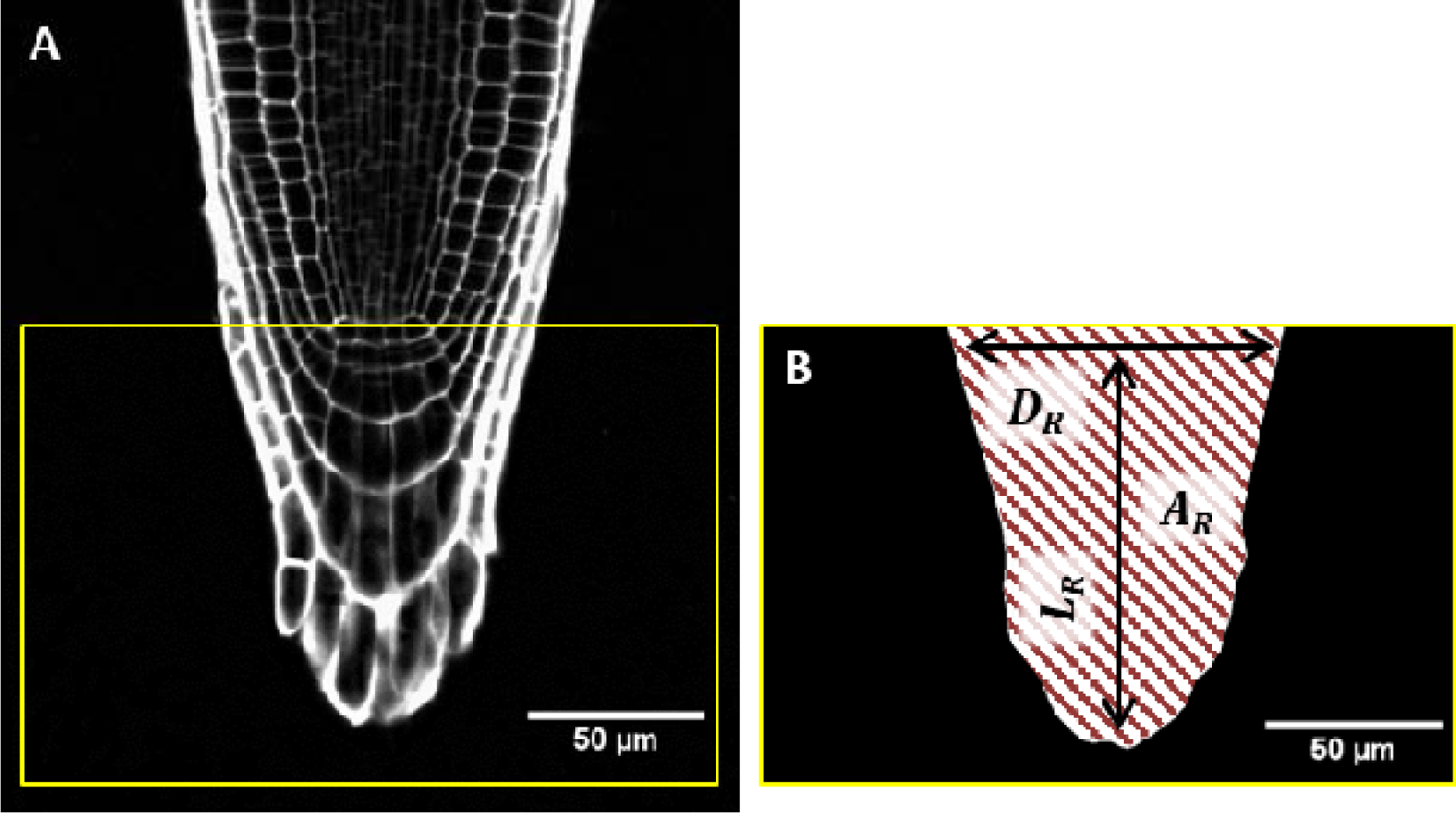
Characterization of the size and shape of the *Arabidopsis thaliana* root tip. (A) Laser scanning microscope image of a PI-stained wild-type 5-day-old root. The yellow square indicates the root tip section on which the size and shape were measured. (B) Root tip section threshold using Image J software. The morphology of the root end was characterized by the length of the root cap L_R_, (from the QC to the root tip), the diameter of the root (at the QC D_R_) and the area of the root tip section A_R_, corresponding to the hatched area. The three measures enabled calculation of the *inverse* Sharpness Index according to equation 4.

The closer *i*SI is to 0, the more the root tip is pointed. Conversely, the closer *SI* is to 1, the more the root tip has a rectangular shape.

### Growth zone length of *Arabidospis thaliana* primary roots

From longitudinal observations of the PI-dyed 5-day-old primary roots under the laser scanning microscope, the length of the cortical cells was measured from the quiescent centre up to 2 mm. Assuming the roots were in steady state, with a constant cell flux, the local velocity over the root tip (*i.e*. the rate at which a cell moves away from the quiescent centre) was calculated from the cell length measurements (see Silk and Erickson, 1979; Silk, 1992; Liu *et al*., 2013 for details). The velocity profile was used to calculate the growth zone length of the roots, corresponding to the length of the root section from the quiescent centre to the point where the velocity reached 94% of the maximal velocity (Fig.S2).

### Maximal axial force of *Arabidospis thaliana* primary roots

The maximal axial force the roots were able to exert on the two-layer medium interface before reorientation of the root tip was estimated for Col-0, *fez-2* and *smb-3* roots grown in 0.2/0.3 medium. After contact between the root tip and the interface, root tip growth induced the deflection of this interface until the root curved or root tip deviated. The shape of the interface just before root tip reorientation was traced using ImageJ software and the maximal deflection (*d*) of the interface was measured with regard to its initial position (before contact with the root tip). The shape of the root tip extremity was fitted by a circle of radius *R*. The maximal force before root reorientation *F*_*crit*_ applied on the surface by the root tip was then estimated using the theory of elastic contact between a sphere and a half-space medium (Hanaor *et al*., 2015),

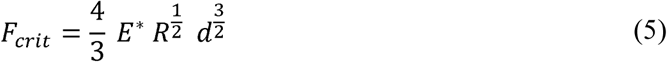

where

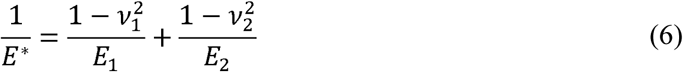

and *E1, E2* are the elastic moduli of the lower medium layer and of the root tip respectively, and *ν*_*1*_ and *ν*_*2*_ are the Poisson ratios of the lower layer and the root tip respectively. Assuming that *E*_*2*_ was much larger than *E*_*1*_, the second part of the equation was neglected while the value of *ν*_*1*_ was estimate at 0.5 (Ahearne *et al*., 2005).

### Time lapse photography and image processing

Root growth and orientation were time-lapsed 5 days after sowing. Two Nikon D7100 cameras equipped with macro lenses (AF-S micro NIKKOR 60mm f/2.8G ED) and controlled by DigiCamControl software, were placed on a rack in front of the culture chambers. Images of the roots growing in the two-layer media were recorded every 5 min for at least 24 hours. All photographs were pre-treated using Rawtherapee software (v4.2.1) and were analyzed using the RootStem Extractor software developed by the PIAF laboratory (Chauvet *et al*., 2016). Briefly, the software enabled extraction of the topological skeleton of each root by screening the pictures perpendicularly to the root axis with a step of 0.3 pixels. The length of the root on each picture and the local inclination as a function of the curvilinear abscissa can be computed from the coordinates of the skeleton. The root length, as a function of time, enabled us to calculate root growth rate by linear fit. Curvature intensity along the root was obtained at each point by deriving the local inclination as a function of time and space. For each root, the curvature intensity data are presented in a graph composed of 3 dimensions: a vertical axis corresponding to root length from the beginning of the measurement, a horizontal axis corresponding to time and a colour bar that depicts curvature intensity. On the spatiotemporal graphs, the green colour points to a straight section of the root, while the red and blue colours depict the highest positive and negative curvatures respectively.

### Gravitropic response of *Arabidospis thaliana* primary roots

*Arabidopsis thaliana* Col-0, *fez-2* and *smb-3* root seedlings were grown vertically during five days in culture chambers containing a 0.5 % Phytagel one-layer medium. The culture chambers were tilted at 90°-*i.e*. horizontal - before recording images of the primary roots every 10 minutes for 22 hours. Analysis of the time-lapse images *via* RootStem Extractor was used to measure root length and root tip angle over time (Fig. S3). These data were used to calculate a gravitropic response, applying the equation described by Chauvet *et al*. (2016).

### Statistical analysis

All data were statistically analyzed using R software (Rstudio, R.3.3.4, https://www.r-project.org/). The means ± standard deviation (std) (or standard error when specified) are presented in the results. The Shapiro-Wilk test was used to test for normal distribution of the data and the Bartlett test was used to test for homoscedasticity. One-way Analysis of Variance (ANOVA) or two-way ANOVA were applied on the data, to determine whether the effect of each factor was significant. When significant differences were observed, post-hoc analyses based on the Tukey test were applied on the data.

## Results

### Phytagel concentration in the medium impacts both medium rigidity and penetration resistance

The elastic modulus E, obtained from the compressive tests, was measured for Phytagel concentrations ranging from 0.2 to 1.2% (Fig. 3A, B). The 0.2%, 0.3%, 0.5% and 1.2% Phytagel growth media showed E values of 3.1 ± 0.2 kPa, 7.6 ± 1.4 kPa, 31.9 ± 6.6 kPa and 158.3 ± 18.3 kPa, respectively (Fig. 3B). The data showed that E was highly dependent on the Phytagel concentration.

**Fig. 3.**
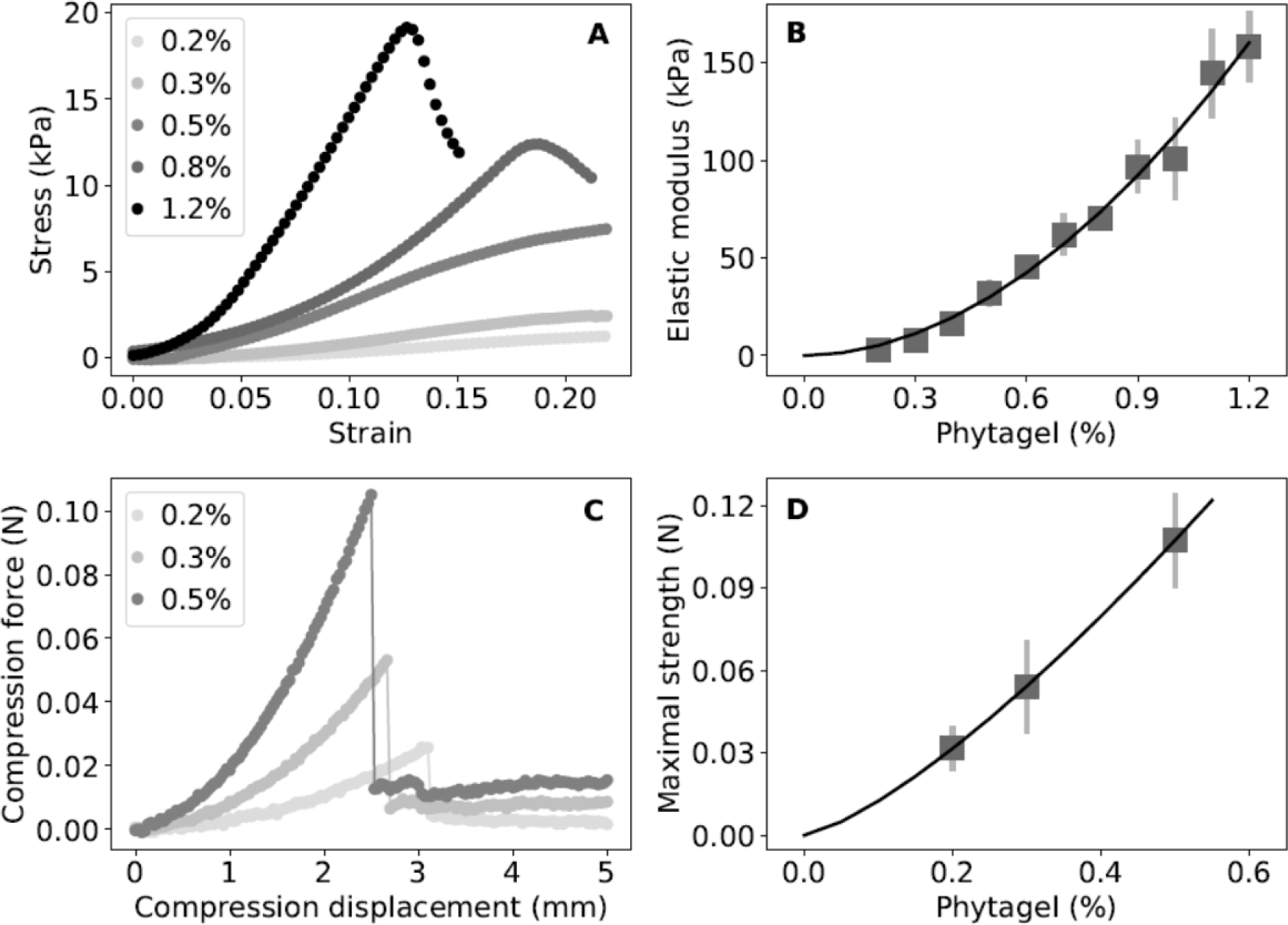
Mechanical properties of Phytagel growth media. (A) Stress-strain curves resulting from compressive tests performed on medium cubes of Phytagel of concentration ranging from 0.2% to 1.2%. The curves correspond to single measurements. (B) The Elastic modulus E was estimated using the slope of the first linear portion of the stress-strain curve. Means ± std are provided, n=3. (C-D) Penetrometer tests performed to investigate the mechanical resistance of the interface in two-layer media. (C) Compression force during penetration of the needle in the lower medium layer of 0.2%, 0.3% and 0.5% Phytagel. The reference starting point for displacement corresponds to the first contact between the needle tip and the interface. The data shown correspond to single measurements. (D) Maximum strength required for the needle to induce rupture of the interface as a function of the Phytagel concentration of the lower medium layer. Means ± std are provided, n=4.

Penetrometer tests were performed in 0.2/0.2, 0.2/0.3 and 0.2/0.5 two-layer media (Fig. 3C, D). The curves of the compression force vs. needle displacement were divided into two steps (Fig. 3C): (I) after the first contact with the lower layer surface, the compression force increased markedly until reaching a maximal value; and (II) while the needle penetrated the lower layer, the loading force rapidly collapsed. The specific pattern of the displacement-force curves illustrated the presence of the interface between the two medium layers, the rupture of which required a specific force, called maximal strength. Interface rupture was induced by a force of 0.03 ± 0.01 N in the 0.2/0.2 medium, 0.05 ± 0.02 N in the 0.2/0.3 medium and 0.11 ± 0.02 N in the 0.2/0.5 medium (Fig. 3D). Thus, the maximal strength increased significantly with the lower medium layer Phytagel concentration. These results show that the maximal strength of the two-layer interface increased with the Phytagel concentration of the lower layer.

### *fez-2* and *smb-3* mutations impact root cap size and shape in *Arabidopsis thaliana* primary roots

Several morphological characterizations were conducted on wild-type (Col-0), *fez-2* and *smb-3* primary roots in order to specify the impact of the mutations on root tip size and shape (Fig. 4). Measures conducted on the primary root extremity of the three *Arabidopsis* lines showed that *fez-2* root cap was shorter and narrower at the base, whereas *smb-3* root cap was longer and larger at the base compared to the wild-type root cap (Fig. 4D, Fig. 4E). However, mean root diameter measured at 500 µm from the root tip, was not significantly different between *fez-2, smb-3* and wild-type roots (Fig. S4). Estimation of the *inverse* Sharpness Index (iSI), of 0.30 ± 0.07 in *fez-2* roots and of 0.41 ± 0.06 in wild-type roots, indicated that *fez-2* root tips were significantly more pointed than wild-type root tips (Fig. 4F). On the contrary, the iSI of the *smb-3* root tips was around 0.5 ± 0.13, illustrating a more rectangular shape (Fig. 4F). Our morphological characterizations indicated that *fez-2* and *smb-3* mutations altered the root cap size and shape in *Arabidopsis thaliana* primary roots. Moreover, the distinct root cap structure of the two mutant lines will make it possible to study the role of root cap shape in root responses to medium strength.

**Fig. 4.**
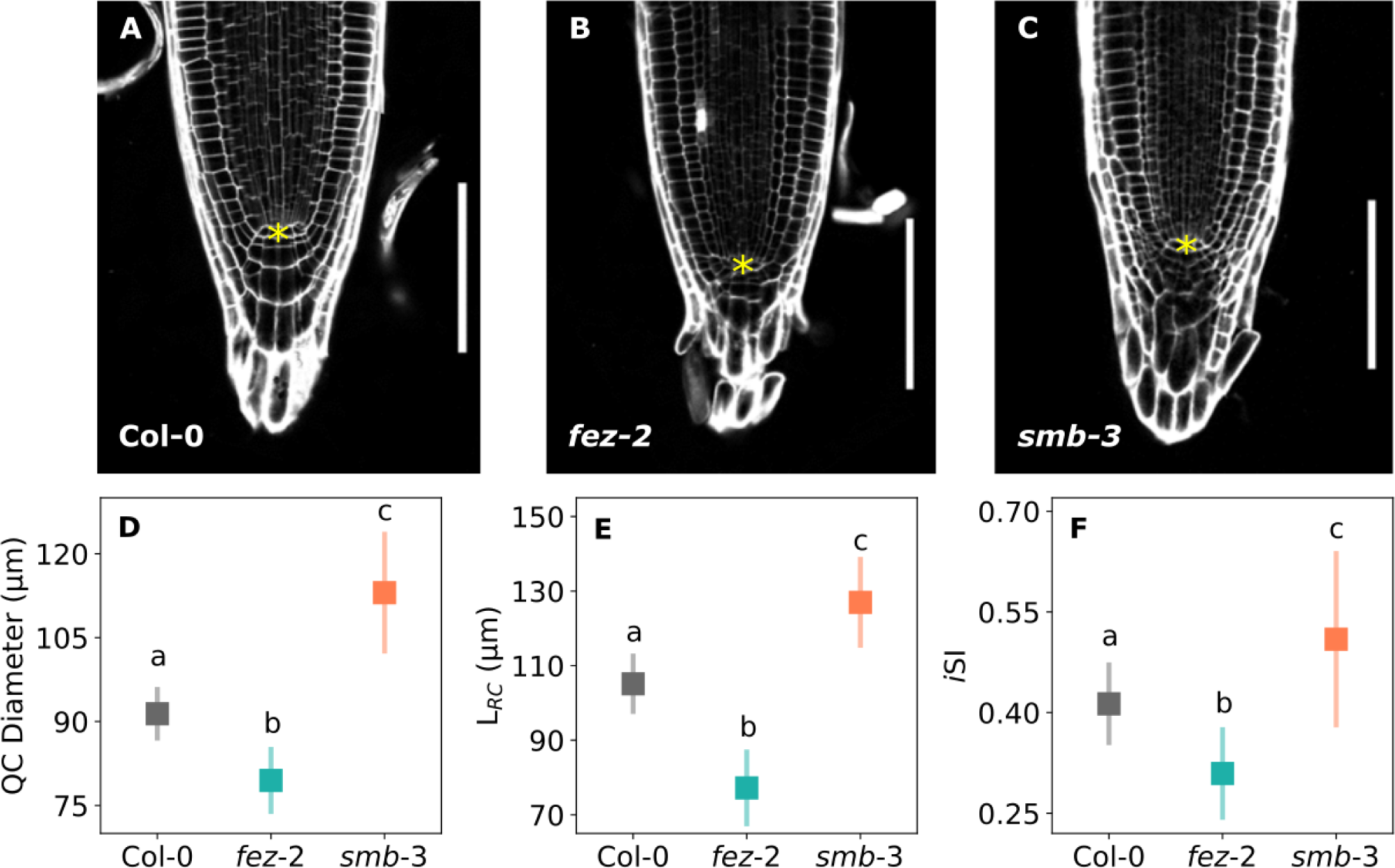
Morphology of wild-type (Col-0), *fez-2* and *smb-3 Arabidopsis thaliana* root tips. (A-C) Representative laser scanning microscope images of Ps-PI stained Col-0 (A), *fez-2* (B) and *smb-3* (C) roots which grew into a compliant growth medium containing 0.2% of Phytagel for five days. Asterisks indicate the quiescent centre. White bar = 100 µm. (D) Diameter of Col-0, *fez-2* and *smb-*3 roots at the quiescent centre. (E) Length of the Col-0, *fez-2*, and *smb-3* root cap L_*RC*_ from the quiescent centre to the root tip. (F) *i*SI of Col-0, *fez-2, smb-3* root tips. (D-F) Means ± std are provided, n= 20. Letters indicate significant differences of means with P<0.05 (ANOVA with Tukey test)

### *smb-3* mutation impacts root growth and gravitropic response whereas *fez-2* mutation does not

From a kinematic analysis, the mean growth zone length, from the quiescent centre to the end of the elongation zone, was estimated at 917 ± 194 µm, 846 ± 303 µm and 898 ± 299 µm in wild-type, *fez-2* and *smb-3* lines, respectively (Fig. 5A, Fig. S2). These similarities suggest that the *fez-2* and *smb-3* mutations may not affect the size of the root growth zone. Analysis of root growth in a 0.2% one-layer Phytagel growth medium showed that the mean growth rate of wild-type (3.2 ± 0.8 µm.min^-1^) and *fez-2* (3.0 ± 0.7 µm.min^-1^) roots was not significantly different (Fig. 5B), whereas *smb-3* roots grew slower, at a mean rate of 1.6 ± 0.3 µm.min^-1^ (Fig. 5B). In the same way, the wild-type and the *fez-2* roots exhibited a similar gravitropic response, of 0.17 ± 0.05 for the wild-type roots and of 0.17 ± 0.02 for the *fez-2* roots, whereas *smb-3* roots exhibited a significantly higher gravitropic response, of 0.36 ± 0.15 (Fig. 5C, Fig. S3). Our studies showed that the *smb-3* mutation impacted the growth and the gravitropic response of the roots, to the contrary of *fez-2* mutation.

**Fig. 5.**
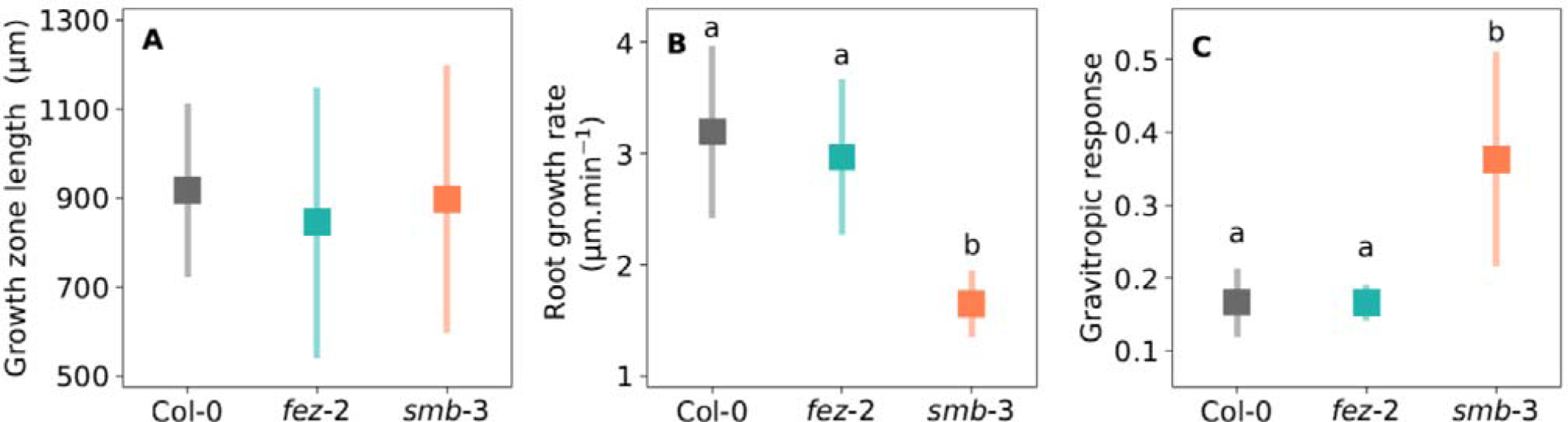
Growth and gravitropic response of wild-type (Col-0), *fez-2* and *smb-3 Arabidopsis thaliana* root tips. (A) Length of the growth zone from the QC to the end of the elongation zone, estimated using a kinematic analysis performed on Col-0, *fez-2* and *smb-3* roots. Means ± std are provided, 14 ≤ n ≤ 18 (B) Root growth rate of Col-0, *fez-2, smb-3* roots growing in compliant one-layer 0.2% Phytagel medium. Means ± std are provided, n= 15. (C) Gravitropic response of Col-0, *fez-2* and *smb-3* roots after tilting at 90°. Means ± std are provided, 14 ≤ n ≤ 16.

### Penetration in harder medium layer is impaired in *Arabidopsis thaliana fez-2* primary roots and enhanced in *smb-3* roots

To understand the role of root cap structure in the root response to an increase in medium strength, the penetration abilities of *fez-2* and *smb-3* primary roots were compared to those of wild-type roots in 0.2/0.2, 0.2/0.3 and 0.2/0.5 two-layer media (Table 1). On the one hand, in the 0.2/0.2 medium, where the increase in medium strength was the weakest, 100% of the wild-type and *smb-3* primary roots progressed into the 0.2% upper layer and crossed the interface to penetrate the 0.2% lower layer. On the other hand, only 83% of the *fez-2* roots penetrated the 0.2% lower layer. In the 0.2/0.3 medium, in response to an intermediate increase in medium strength, 60% of the *Arabidopsis thaliana* primary roots penetrated the lower layer. Root penetration into the 0.3% Phytagel lower layer was also observed for 28% of the *fez-2* primary roots and 80% of the *smb-3* primary roots. Finally, in the 0.2/0.5 medium, only 1.7 % of the wild type primary roots and 18.3% of the *smb-3* roots penetrated the 0.5% lower layer. None of the *fez-2* roots penetrated the lower layer in response to the most important increase in medium strength.

**Table 1.** Penetration ability of *Arabidopsis thaliana* wild-type (Col-O), *fez-2*, and *smb-3* primary roots in two-layer media. Penetration ability is represented by the percentage of roots that were able to penetrate 0.2%, 0.3% and 0.5% lower layers. For each genotype and condition, 6 replicates of 10 roots were observed 8 days after germination. The values indicate the average percentage over the six replicates ± std. The letters show significant difference of means with P<0.05 (two-way ANOVA with Tukey test)

| Two-layer medium | Penetrating roots (%) |  |  |
| --- | --- | --- | --- |
|  | 0.2/0.2 | 0.2/0.3 | 0.2/0.5 |
| Col-0 | 100.00 ± 0.00 <sup>a</sup> | 60.00 ± 3.65 <sup>c</sup> | 1.67 ± 1.67 <sup>e</sup> |
| *fez-2* | 83.33 ± 4.94 <sup>b</sup> | 28.33 ± 3.07 <sup>d</sup> | 0.00 ± 0.00 <sup>e</sup> |
| *smb-3* | 100.00 ± 0.00 <sup>a</sup> | 80.00 ± 5.16 <sup>b</sup> | 18.33 ± 4.77 <sup>d</sup> |

These results showed that the *fez-2* roots exhibited impaired penetration ability in response to an increase in medium strength, whereas the *smb-3* roots exhibited enhanced penetration ability compared to the wild type roots. To understand where these differences in root penetration ratio come from, the critical axial force that each root was able to apply on the two-layer medium interface before tip reorientation was estimated.

### Critical axial force is impaired in *Arabidopsis thaliana fez-2* roots and enhanced in *smb-3* roots

The critical axial force of wild-type, *fez-2* and *smb-3* primary roots was estimated for the roots growing in the 0.2/0.3 medium, in which the greatest differences in penetration ratio were observed between the three lines (Fig. 6). In the 0.2/0.3 medium, the roots grew into the upper layer before encountering the interface between the two layers. After contact with the interface, root axial growth induced deflection of the interface until either the rupture of the interface or initiation of a zone of curvature. Deflection of the interface *d* before curvature initiation was measured for each wild-type, *fez-2* and *smb-3* root which did not penetrate the lower layer (Fig. 6A). Wild-type roots displaced the interface at a mean maximal depth *d* of 0.37 ± 0.03 mm, the *fez-2* roots displaced it at a mean *d* of 0.27 ± 0.01 mm and the *smb-3* roots displaced it at a mean *d* of 0.49 ± 0.03 mm (Fig. 6B). The values of *d*, together with the radius of the root tips and the rigidity of the lower medium layer (E = 7.6 kPa) made it possible to estimate the critical axial growth force of wild-type, *fez-2* and *smb-3* primary roots (Fig. 6C). Mean critical force was estimated at 1.12 ± 0.14 10^-3^ N for wild-type roots, 0.66 ± 0.07 10^-3^ N for *fez-2* roots and 1.77 ± 0.18 10^-3^ N for *smb-3* roots, confirming that *smb-3* roots were able to apply a higher axial force on the interface while *fez-2* roots applied a weaker axial force than wild type roots (Fig. 6C).

**Fig. 6.**
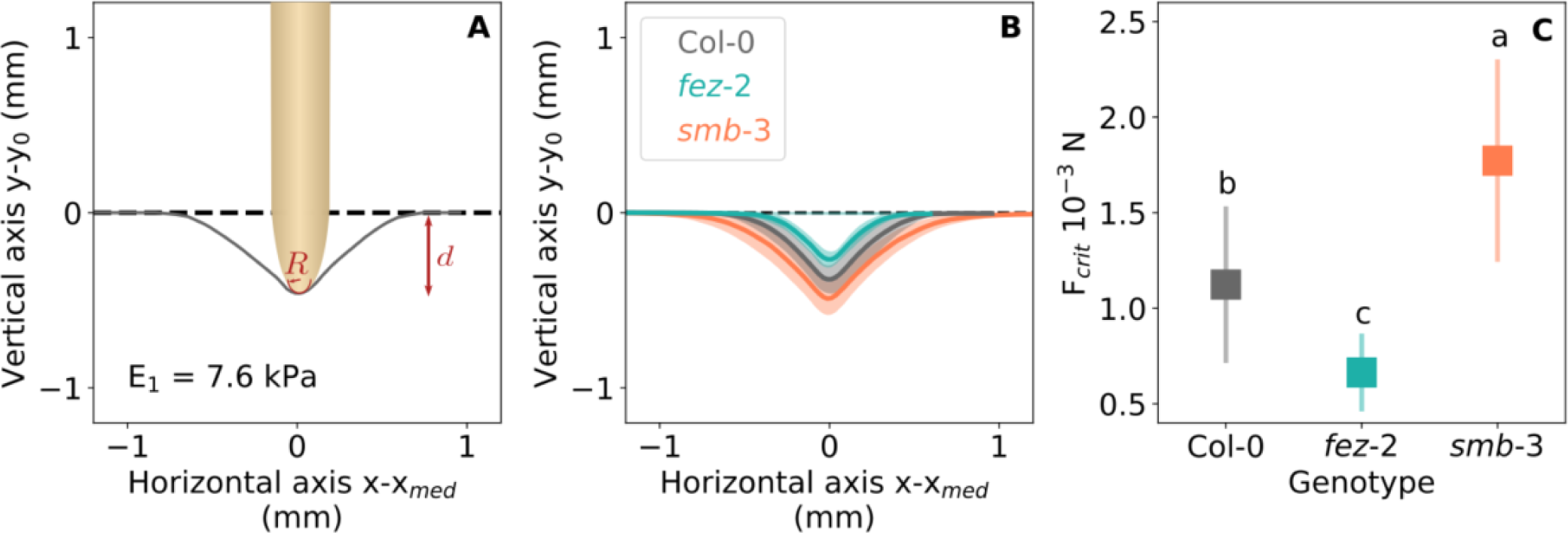
Critical axial force of wild-type (Col-0), *fez-2* and *smb-3 Arabidopsis thaliana* root. (A) Schematic example of a root tip growing against the 0.2/0.3 interface of elastic modulus E_1_= 7.6 kPa. The grey curve represents the shape of the interface just before root curvature initiation. *d* is the corresponding maximal deflection of the interface. R is the radius of the root tip in contact with the surface. The dashed line corresponds to the initial shape of the interface before contact with the root tip. (B) Shapes of the interface (E = 7.6 kPa) deformed by Col-0, *fez-2* and *smb-3* root tips just before their curvature initiation. Lines are means, n=10. The dashed line corresponds to the initial shape of the interface before contact with the root tip. (C) Evaluation of the critical force Fcrit that the Col-0, *fez-*2 and *smb-3* roots were able to apply on the interface before reorienting. Fcrit was computed according to the equation 5. (n=10). Means ± std are provided. The letters indicate significant difference of means with P<0.05 (ANOVA with Tukey test).

### The *Arabidopsis thaliana fez-2* and *smb-3* roots exhibit altered curvature establishment in response to an increase in medium strength

In two-layer media, roots that did not apply a sufficient axial force to induce rupture of the interface reoriented their growth thanks to the initiation of zones of curvature. The initiation of zones of curvature after contact with the interface was spatiotemporally studied as a function of the increase in medium strength and as a function of the genotype (Fig. 7-9, Fig. S5-S11). Representative examples of wild-type, *fez-2* and *smb-3* roots reorienting their growth in 0.2/0.3 and 0.2/0.5 media are presented on Fig. 7-9.

**Fig. 7.**
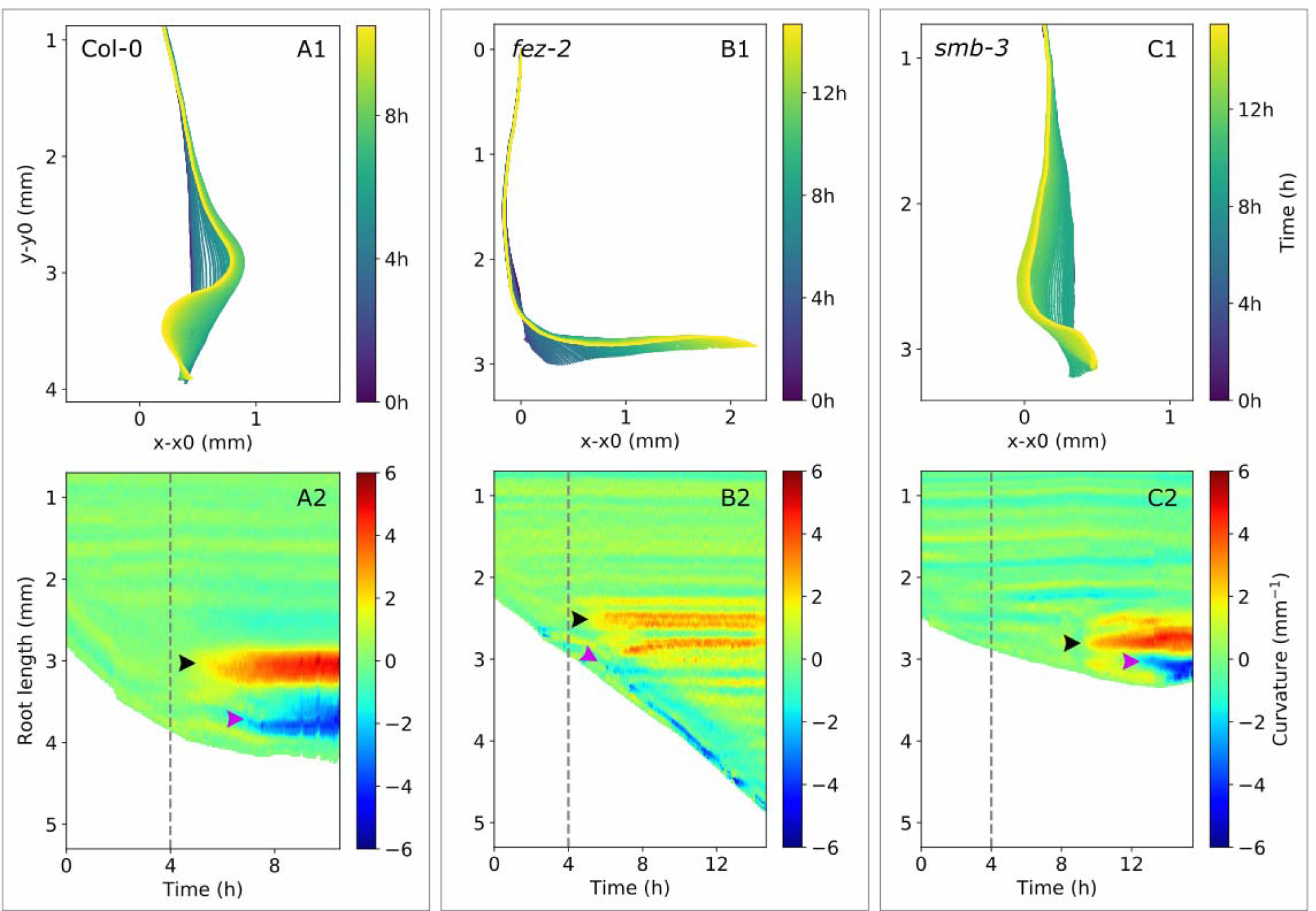
Spatiotemporal reorientation of wild-type (Col-0), *fez-2* and *smb-3 Arabidopsis thaliana* root tips in the 02/0.3 medium. (A) Skeleton (A1) and spatiotemporal analysis (A2) of a Col-0 primary root reorienting after contact with the 0.2/0.3 interface. (B) Skeleton (B1) and spatiotemporal analysis (B2) of a *fez-2* primary root reorienting after contact with the 0.2/0.3 interface. (C) Skeleton (C1) and spatiotemporal analysis (C2) of a *smb-3* primary root reorienting after contact with the 0.2/0.3 interface. In the spatiotemporal graphs, root length as a function of time is presented. The colours represent curvature intensity: negative for right-handed curvatures and positive for left-handed curvatures. The green colour corresponds to the straight zones of the roots, while the red and blue colours depict the highest positive and negative curvatures respectively. The dashed lines indicate the contact between the root tip and the interface. Black and purple arrowheads indicate the initiation of the first and the second zone of curvature, respectively.

Following contact with the 0.3% interface, non-penetrating roots firstly grew downward against the interface and deformed it until a first zone of curvature was formed (Fig. 7). This first zone of curvature appeared at 515 ± 98 µm, 603 ± 33 µm and 398 ± 53 µm from the root tip extremity in wild-type, *fez-2* and *smb-3* roots, respectively. Once established, the curvature covered a zone of 657 ± 121 µm, 977 ± 58 µm and 414 ± 73 µm in wild-type, *fez-2* and *smb-3* roots, respectively (Fig. 7, Table S1). The length of the zone of curvature was significantly greater in *fez-2* roots than in *smb-3* roots but no significant differences were observed between wild type and mutant roots (Table S1). In the 0.2/0.3 medium, initiation of the first zone of curvature induced lateral deviation of *fez-2* root tips whereas wild-type and *smb-3* root tips continued to grow downwards. These distinct behaviours led to the establishment of two distinct root shapes (Fig. 8, Fig. S5-7). In wild type and *smb-3* roots, several zones of curvature were initiated successively and their establishment shared common characteristics, leading to the formation of one or more curls (Fig. 8A., Fig. S5-6). Thus, in the 0.2/0.3 medium, wild-type and *smb-3* roots developed a “curly” shape. In *fez-2* roots, initiation of the first zone of curvature and subsequent root tip deviation were followed by the initiation of a second zone of curvature, the characteristics of which were distinct from those of the first curvature (Fig. 7B2, Fig. 8B). In the 0.2/0.3 medium, establishment of the two distinct zones of curvature in *fez-2* roots led to the formation of a “step-like” shape (Fig. 8B). The second zone of curvature was formed at 280 ± 23 µm from the root tip extremity and extends over 455 ± 45 µm (Fig. 7B2). Moreover, whereas the first zone of curvature was maintained after its establishment, the second one moved and kept a constant distance from the root tip extremity (Fig. 7).

**Fig. 8.**
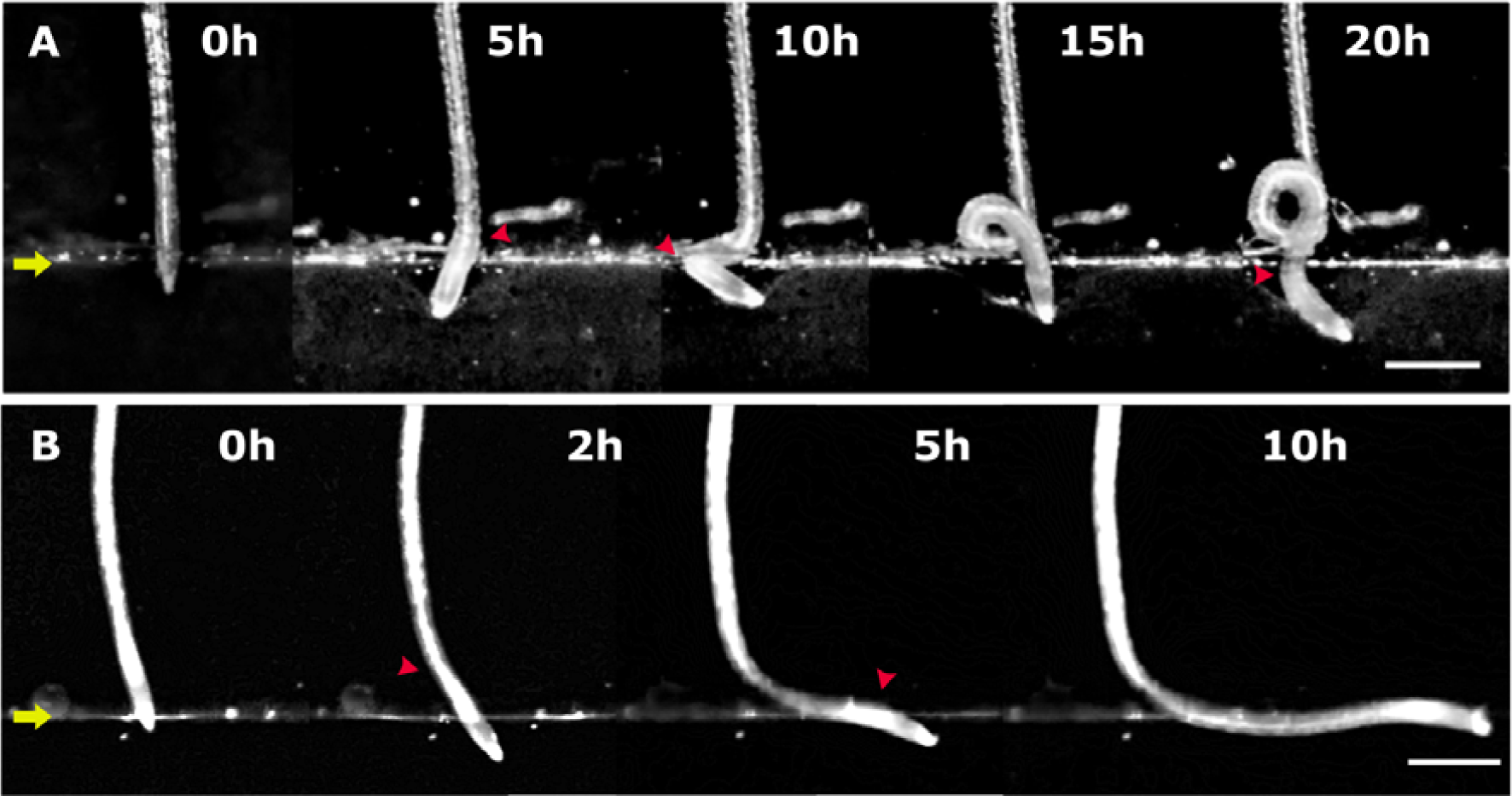
Major time-steps of the establishment of the curly (A) and step-like (B) root morphologies after the contact between the root tip and the interface of the 0.2/0.3 medium. (A) Root tip reorientation and curvature formation of a representative wild-type root which did not penetrate the lower layer in the 0.2/0.3 medium, at 0 h, 5 h, 10 h, 15 h and 20 h after contact with the interface. This root developed a characteristic curly morphology. (B) Root tip reorientation and curvature formation of a representative *fez-2* root that did not penetrate the lower layer in the 0.2/0.3 medium, at 0 h, 2 h, 5 h and 10 h after contact with the interface. This root developed a characteristic step-like morphology. Yellow arrows indicate the 0.2/0.3 interface. Red arrowheads indicate the formation of zones of curvature. White bars correspond to 500 µm

In the 0.2/0.5 medium, all the wild-type and *fez-2* roots developed a step-like shape following contact with the interface (Fig. 9, Fig. S8-9). For the *smb-3* line, three distinct root shapes were observed in the 0.2/0.5 medium: “curly”, “wavy” and “step-like” shapes (Fig. S10-11). The spatiotemporal establishment of the step-like shape of wild-type, *fez-2* and *smb-3* primary roots facing the most important increase in medium strength was compared. After contact with the 0.2/0.5 interface, a first zone of curvature was formed at 540 ± 50 µm, 664 ± 64 µm and 407 ± 32 µm from the root tip extremity in wild-type, *fez-2* and *smb-3* roots, respectively (Fig. 9). In *fez-2* roots, curvature was initiated significantly further from the root tip extremity than in *smb-3* roots but no significant differences were observed between the wild type and the mutant roots (Table S2). Once established, the curvature extended over a zone of 657 ± 121 µm, 977 ± 58 µm and 414 ± 73 µm in wild-type, *fez-2* and *smb-3* roots, respectively (Fig. 9). Thus, the zone of curvature was significantly longer in *fez-2* roots than in wild-type and *smb-3* roots. Moreover, maximal curvature intensity was significantly weaker in *fez-2* roots (2.3 ± 0.5 mm^-1^) than in wild type (4.7 ± 0.5 mm^-1^) and *smb-3* roots (4.8 ± 0.6 mm^-1^) (Fig. 9, Table S2). While the root tip reoriented its growth, all the wild-type, *fez-2* and *smb-3* roots showed a second zone of curvature (Fig. 9). Establishment of this second curvature shared common characteristics between the three Arabidopsis lines, except for the maximal intensity, which was significantly weaker in *fez-2* roots than in wild-type and *smb-3* roots (Fig. 9, Table S2). As was observed in *fez-2* root tips reorienting their growth in the 0.2/0.3 medium, the second zone of curvature remained at a constant distance from the root tip during root growth along the 0.2/0.5 interface (Fig. 9).

**Fig. 9.**
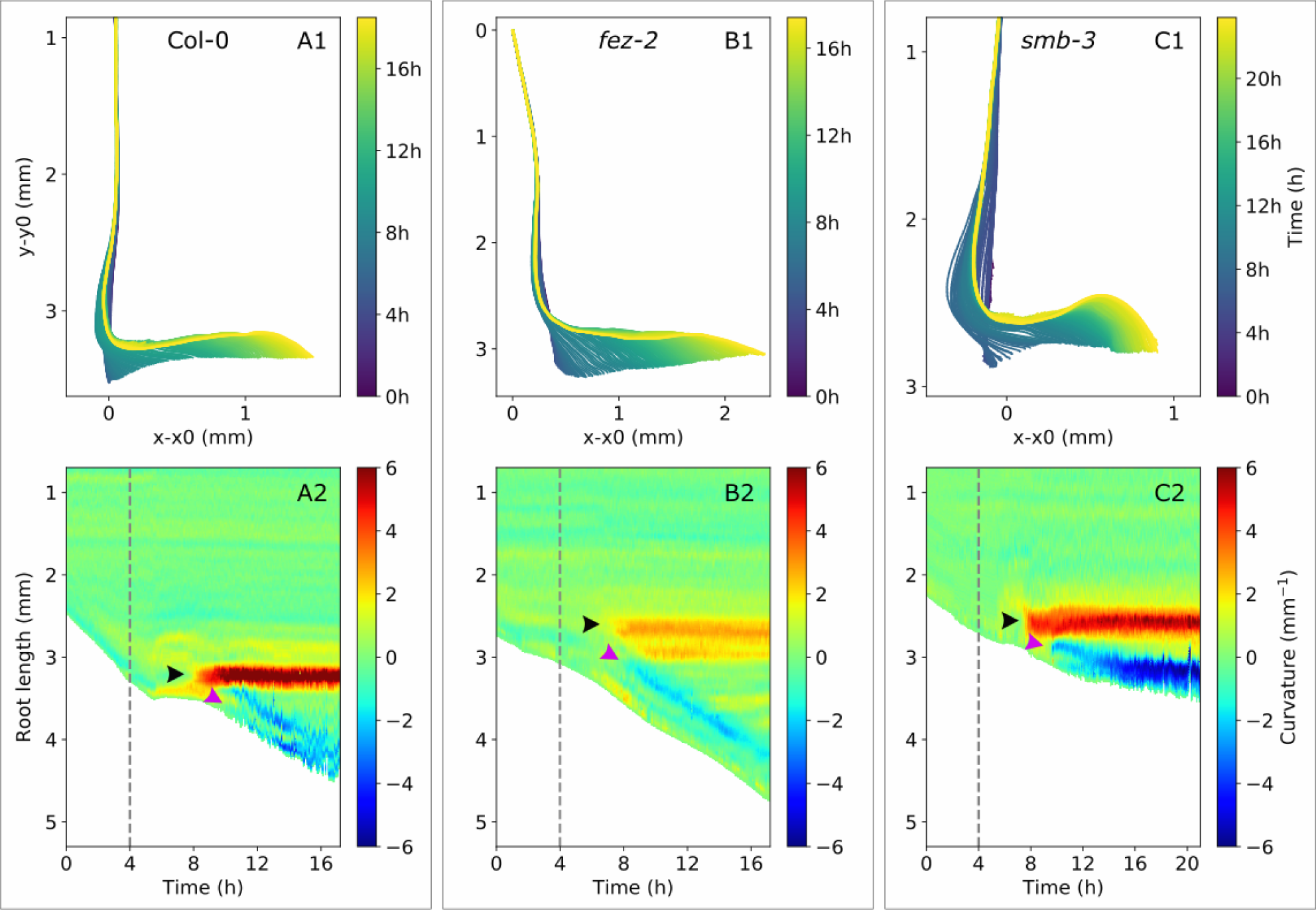
Spatiotemporal reorientation of wild-type (Col-0), *fez-2* and *smb-3 Arabidopsis thaliana* root tips in the 0.2/0.5 medium. (A) Skeleton (A1) and spatiotemporal analysis (A2) of a Col-0 primary root reorienting after contact with the 0.2/0.5 interface. (B) Skeleton (B1) and spatiotemporal analysis (B2) of a *fez-2* primary root reorienting after contact with the 0.2/0.5 interface. (C) Skeleton (C1) and spatiotemporal analysis (C2) of a *smb-3* primary root reorienting after contact with the 0.2/0.5 interface. In the spatiotemporal graphs, root length as a function of time is presented. The colours represent curvature intensity: negative for right-handed curvatures and positive for left-handed curvatures. The green colour corresponds to the straight zones of the roots. The dashed lines indicate contact between the root tip and the interface. Black and purple arrowheads indicate the initiation of the first and the second zone of curvature, respectively.

The two-layer media experiments illustrated that root tip reorientation in response to an increase in medium strength was not only impacted by the mechanical resistance of the interface but also by the size and shape of the root cap.

### The increase in medium strength impacts the root growth rate in wild-type, *fez-2* and *smb-3 Arabidopsis thaliana* primary roots differently

Growth rate was monitored for wild-type, *fez-2* and *smb-3* roots in 0.2/0.2, 0.2/0.3 and 0.2/0.5 media. In the 0.2/0.2 medium, only the growth rate of roots which penetrated the lower layer was measured (Table S3). In the 0.2/0.3 medium, the growth rate was measured for both penetrating and reorienting roots of all three lines (Table S3, Table S4, Fig. 10A). In wild-type reorienting roots, the contact between the root tip and the interface was followed by a significant 60% decrease in mean growth rate (Fig. 10A). Unfortunately, our experimental system based on 2D visualization of the roots prevented measurement of the root growth rate until the end of the root tip reorientation. A significant 60% decrease in mean root growth rate was also observed in *smb-3* reorienting roots whereas no significant decrease was observed in *fez-2* reorienting roots after contact between the root tip and the 0.2/0.3 interface (Fig. 10A, Table S4).

**Fig. 10.**
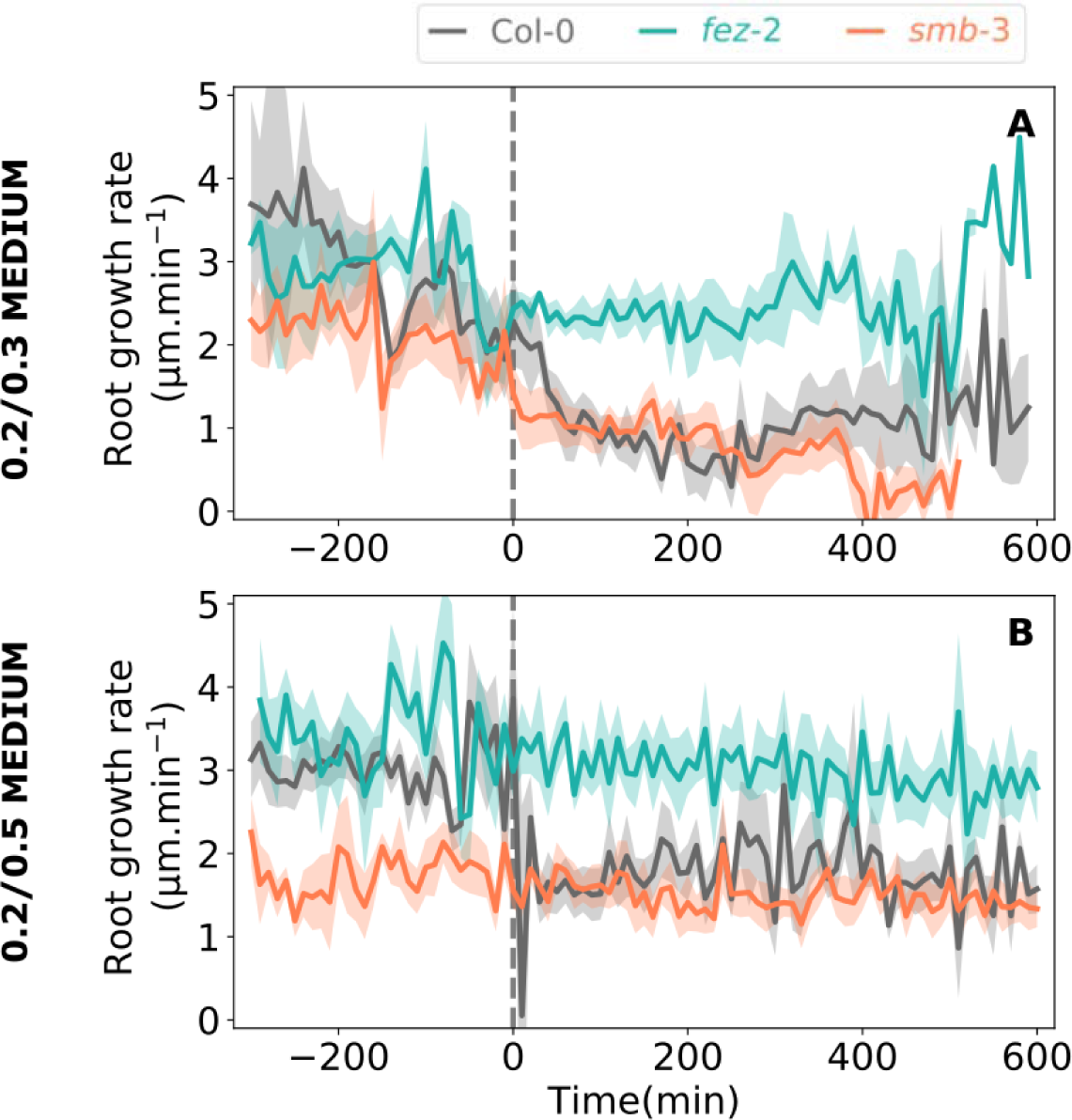
Growth rate of wild-type (Col-0), *fez-2, smb-3 Arabidopsis thaliana* roots in two-layer media before and after contact with the interface. (A-B) Root growth rate of Col-0, *fez-2*, and *smb-3* roots that reoriented their growth after contact with the interface in the 0.2/0.3 medium (A) and in the 0.2/0.5 medium (B). Means ± standard error are provided, 8<n<12. Dashed lines indicate the contact between the root tip and the interface.

In the 0.2/0.5 medium, the growth rate was measured for the reorienting roots from the wild-type, *fez-2* and *smb-3* lines (Fig. 10B). The growth rate of wild-type roots, which was 3.0 ± 0.2 µm.min^-1^ in the 0.2 % lower layer, decreased by 37% to 1.9 ± 0.3 µm.min^-1^ after contact with the interface. When root tip reorientation was complete, the initial growth rate did not recover. After contact with the 0.2/0.5 interface, the establishment of the two zones of curvature in *smb-3* roots was also accompanied by a significant 44% decrease in mean growth rate (Fig. 10B; Table S4). However, the *fez-2* root showed no variation in growth rate after contact with the interface in the 0.2/0.5 medium.

Our results show that the increase in medium strength in the two-layer media impacts the growth rate of wild-type roots. Moreover, this impact seems to be altered in mutant roots, and especially in *fez-2* roots, the growth rate of which does not seem to be affected by contact with the interface.

## Discussion

Our experimental design has proven to be relevant for investigating root responses to medium strength. Precise characterization of media mechanical properties showed strong correlations between medium rigidity, penetration resistance and Phytagel concentrations. Moreover, precise spatiotemporal analysis of root growth rate and root tip orientation allowed us to discriminate the passive and active mechanisms involved in root tip reorientation. Complete characterization of root tip reorientation led to a better understanding of the role of the root cap structure in the root responses to mechanical stresses.

### Root tip reorientation involves both passive and active mechanisms

In our experimental system, roots grew through a first medium layer of low strength until it came across the interface that led to an increase in medium strength. In response to the intermediate and the highest increases in medium strength, wild-type root growth rate decreased significantly. This suggests the existence of a biological process that senses the increase in medium strength and that subsequent responses are triggered in the wild-type roots. This decrease, observed for both penetrating and non-penetrating roots, was comparable to that observed in poplar roots submitted to a physical obstacle (Bizet *et al*., 2016).

Root growth against the interface generates an increasing compression reaction force on the root (Kolb *et al*., 2017). If the compression force reaches a critical value while the root is still in contact with the obstacle, Euler’s buckling can occur (Bizet *et al*., 2016). Buckling was observed for wild-type, *fez-2* and *smb-3* non-penetrating roots and led to three distinct shapes depending on the genotype and on the strength of the interface. When they encounter an intermediate increase in medium strength, non-penetrating roots mostly developed a curly-shape. This shape was already observed in *Medicago truncatula* roots growing in a two-layer Phytagel medium (Silverberg *et al*., 2012). From a mathematical model based on the theory of buckling rods, the authors assigned the curly shape of M*edicago* roots to a combination of helical buckling and simultaneous twisting near the root tip. Lateral root containment by the upper medium layer seemed to be a key feature modulating root helical buckling (Silverberg *et al*., 2012). Poplar roots grown in non-constraining hydroponic solution and submitted to a physical obstacle buckled rapidly with a long wavelength curve shape (Bizet *et al*., 2016). In our experimental system, some *smb-3* non-penetrating roots adopted this shape that we called the “wavy” shape. In the two-layer media, since all the upper layers had the same mechanical strength, the various shapes adopted by roots cannot be attributed to their constriction level but more likely to their variability in anchorage or diameter. For most of the curly-shaped and wavy-shaped roots, the first curvature was initiated in a zone between 400 and 700 µm from the root tip. According to our kinematic analyses performed on roots growing in a compliant one-layer medium, this zone corresponds to the zone where the maximal elongation rate occurs. Thus, the zone of curvature initiation may correspond to the zone of mechanical weakness described in Bizet *et al*. (2016). To confirm this, *in-vivo* kinematic analyses should be carried out on roots growing in two-layer media. Our results, together with previous studies, suggest that curly and wavy shapes are mainly driven by the mechanical response of the root acting as a “growing” rod whose end is blocked by the obstacle.

The step-like shape was developed in roots mostly in response to the highest increase in medium strength. In this case, the first curvature induced deviation of the root tip, which implied a change in mechanical forces applied on the root tips from axial to asymmetric radial forces. The rapidity and position of curvature initiation suggest that it may be interpreted as buckling. The first curvature was followed by the initiation of a second, closer to the root apex, allowing the root tip to regain a vertical position. This second curvature provided the root tip with a steady form from changing cells, and was previously described as the result of the interaction between gravitropic and thigmotropic responses (Massa and Gilroy, 2003). Root tip deviation was preceded by a decrease in root growth rate which did not recover after relaxation of axial mechanical stresses. The initiation of the second curvature and the maintenance of the reduced root growth rate suggest that, even if the first curvature was initially provoked by buckling, the further root growth direction may be reoriented in an active way.

The different patterns of curvature establishment suggest that the mechanical strength of the interface determines the subsequent root response (buckling or bending). A middle-strength interface may allow the root tips to sink into it, preventing their deviation after root buckling. On the contrary, a strong interface seems to be barely deformed by the root tips, which may be free to deviate and may trigger active root reorientation.

### Root cap structure impacts the root responses to an increase in medium strength in the opposite way than that expected

Based on previous studies (Vollsnes *et al*., 2010; Ruiz *et al*., 2016; Colombi *et al*., 2017*b*), we hypothesized that root cap shape influenced root penetration ability and root response to medium strength by changing the distribution of mechanical stresses resulting from the axially-oriented external force. It has been suggested that a pointed root tip increased axial stresses at the tip forefront rather than around the tip, resulting in enhanced crack initiation and enhanced penetration ability (Colombi *et al*., 2017*b*). In our experiments, contrary to this hypothesis, *fez-2* roots, that showed acute tips, exhibited reduced penetration abilities whereas *smb-3* roots, that showed dome-shaped tips, exhibited enhanced penetration abilities compared to wild-type roots. In mutant roots, and especially in *fez-2* roots, since the morphological alterations reside mainly in the structure of the cap, our results suggest that the structure of the root cap have an impact on the establishment of the responses to medium strength but in a way that contradicts our hypothesis. This suggests either that the root cap shape is not the key feature determining the root responses to medium strength or that another factor could influence the establishment of the responses. To go further, attention should be paid to the cap rigidity of wild-type and mutant roots. The loss or the accumulation of root cap cell layers could alter not only the cap shape but also its rigidity. A softer cap would deform more easily under an increase in medium strength, resulting in the modification of its shape. In *fez-2* roots for example, the benefit of an acute tip shape on penetration abilities could be minimized by low root cap rigidity.

The difference in penetration abilities between mutant and wild-type roots may be assigned to their different critical buckling forces. The *fez-2* non-penetrating roots mostly exhibited a step-like shape even when they experienced the weakest increase in medium strength. This suggests that the low resistance of *fez-2* roots to buckling did not allow them to apply the mechanical stress that is required to break the interface, while it did not prevent root tip deviation. On the contrary, the higher buckling resistance of *smb-3* roots allowed them to apply more axial stress on the lower layer surface and thus to penetrate the strongest layers more easily. The differences in root critical axial force between the three *Arabidopsis* lines may be explained either by differences in diameter or rigidity at the buckling root zone, or by differences in growth zone length, as defined by Euler’s law. Our characterizations of the mutant root morphologies did not reveal any alteration in root diameter or growth zone length compared to the wild-type roots in low medium-strength conditions. Our results suggest that either the rigidity of mutant roots was altered compared to that of wild-type roots or that growth zone length, tissue rigidity and/or root diameter may be modulated differently in response to an increase in medium strength between the three *Arabidopsis* lines.

Further experiments should be conducted to confirm either of these hypotheses, such as AFM assays on wild-type and mutant roots in order to accurately characterize root tissue local rigidity. Moreover, kinematic *in-vivo* analyses could be performed on roots growing in the two-layer media in order to define the impact of the increase in medium strength on the growth zone length or root cell elongation. In the meantime, if the second hypothesis turns out to be true, we may speculate that the increase in medium strength is perceived by root cap cells and that signal transduction triggers root responses in the elongation zone. In *fez-2* roots, the loss of cap cell layers may alter the mechanosensitivity and thus the establishment of root responses to an increase in medium strength. The fact that the growth rate of *fez-2* roots was not affected by medium strength consolidates this hypothesis. A properly formed root cap may be required to trigger the signal perception and transduction processes within the root and thus to allow the root to respond to an increase in medium strength.

## Conclusion

During their progression in soil, roots frequently face physical heterogeneities they have to move or cross by generating mechanical forces or circumvent by reorienting their growth. Our experimental approach provides new insights into the role of the root cap on root penetration and reorientation in response to variations in medium strength. When Arabidopsis primary roots meet a new medium that shows higher mechanical resistance, they adopted three main types of responses: root penetration in a brittle medium, or root buckling without tip deviation leading to a curly shape in a middle-strength medium, or root buckling followed by tip deviation leading to a step-like shape in the strongest medium. The responses to variation in medium strength were altered in *fez-2* and *smb-3* mutant roots, which exhibit a more acute and more rectangular tip shape, respectively. The alteration in penetration ability, reduced in *fez-2* roots and enhanced in *smb-3* roots, seems to result from altered resistances to buckling. These results suggest that the structure of the root cap could impact root responses to medium strength by influencing root resistance to buckling.

## Supplementary data

Fig. S1. The two-layer Phytagel media 0.2/0.2, 0.2/0.3 and 0.2/0.5 were used to study the responses of *Arabidopsis thaliana* roots to an increase in medium strength.

Fig. S2. Cell length and velocity profiles of wild-type (Col-0), *fez-2* and *smb-3 Arabidopsis thaliana* primary roots.

Fig. S3. Gravitropic response of wild-type (Col-0), *fez-2* and *smb-3* primary roots.

Fig. S4. Diameter of the *Arabidopsis thaliana* primary roots of the wild-type (Col-0), *fez-2* and *smb-3* lines at 500 µm from the root tip extremity.

Fig. S5. Time-lapse movie of a reorienting wild-type root in the 0.2/0.3 two-layer medium Fig. S6. Time-lapse movie of a reorienting *smb-3* root in the 0.2/0.3 two-layer medium Fig. S7. Time-lapse movie of a reorienting *fez-2* root in the 0.2/0.3 two-layer medium

Fig. S8. Time-lapse movie of a reorienting wild-type root in the 0.2/0.5 two-layer medium

Fig. S9. Time-lapse movie of a reorienting *fez-2* root in the 0.2/0.5 two-layer medium

Fig. S10. Time-lapse movie of a reorienting step-like shape *smb-3* root in the 0.2/0.5 two-layer medium

Fig. S11. Time-lapse movie of a reorienting wavy shape *smb-3* root in the 0.2/0.5 two-layer medium

Table S1. Characteristics of the initiation of zones of curvature after contact with the interface in wild-type (Col-0), *fez-2* and *smb-3* primary roots which did not penetrate the lower layer in the 0.2/0.3 medium.

Table S2. Characteristics of the initiation of zones of curvature after contact with the interface in wild-type (Col-0), *fez-2* and *smb-3* primary roots which did not penetrate the lower layer in the 0.2/0.5 medium.

Table S3. Detailed growth rate of penetrating roots of the wild-type (Col-0), *fez-2* and *smb-3* Arabidopsis lines in the 0.2/0.2 and 0.2/0.3 two-layer media

Table S4. Detailed growth rate of reorienting roots of the wild-type (Col-0), *fez-2* and *smb-3* Arabidopsis lines in the 0.2/03 and 0.2/0.5 two-layer media.

## Acknowledgements

We would like to thank Ben Scheres and Viola Willemsen (University of Utrecht, The Netherlands) for the kind mutant line donations. This research received financial support from the French Space Agency (CNES) and by the recipient of INRA programs. J.R. received a doctoral grant from the French National Ministry of Education and Research. H.C received a Postdoctoral grant from the Auvergne-Rhône Alpes Council with cofunding from FEDER. F. B. received a Postdoctoral grant from the CNES.

